# rtmsEcho: An Open-Source R Package for Automated Analysis of Acoustic Ejection Mass Spectrometry Data

**DOI:** 10.1101/2025.06.05.658081

**Authors:** Mary Ashley Rimmer, Nathaniel Twarog, Tharindu Ranathunge, Jingheng Wang, Yong Li, Taosheng Chen, Anang Shelat, Lei Yang

## Abstract

Mass spectrometry (MS) is a well-established technology in biological research, enabling sensitive and precise quantitative analysis of complex samples. While traditional LC-MS systems provide robust performance for targeted analyses, their reliance on chromatographic separation limits throughput, rendering large-scale studies inefficient. The emergence of acoustic ejection mass spectrometry (AEMS) has revolutionized high-throughput workflows by eliminating chromatography and enabling direct nanoliter-scale sampling, achieving hundreds to thousands of measurements per hour. However, AEMS’s full potential remains constrained by software limitations—existing tools lack robust automated processing capabilities for critical tasks such as peak detection, integration, and multimodal data analysis (e.g., multiple reaction monitoring, precursor ion, and neutral loss scans). To address this gap, we developed rtmsEcho, an open-source R package that extends our previously published rtms framework. This specialized solution provides direct access to AEMS data, enabling customizable processing of both MRM and full-scan acquisitions (precursor ion and neutral loss modes) while automating shot-to-peak association and spectral analysis. By streamlining data extraction and quantification, rtmsEcho enhances efficiency and reproducibility in high-throughput applications, including drug discovery, quality control, and clinical diagnostics. This innovation bridges a critical gap in AEMS data analysis, allowing researchers to fully leverage the speed and precision of next-generation mass spectrometry.

## INTRODUCTION

Mass spectrometry (MS) has emerged as an indispensable analytical technique in pharmaceutical,^1^ biotechnology, clinical,^2^ and research laboratories,^3, 4^ enabling sensitive and quantitative analysis of small molecules and peptides in complex matrices.^5^ With applications spanning drug discovery, biomarker validation, and systems biology, MS provides the accurate quantification required for critical screening and quality control application.^6-8^ The current gold standard for targeted analysis employs liquid chromatography (LC) coupled with mass spectrometry (MS), typically using single quadrupole (Q) or triple quadrupole (QQQ) instruments operating in single-ion monitoring (SIM), selected reaction monitoring (SRM), or multiple reaction monitoring (MRM) modes due to their superior selectivity, sensitivity, and robustness.^9, 10^ However, the intrinsic constraints of chromatographic separation, such as mandatory equilibration times and instrument cycle requirements, cause throughput limitations. Even with highly optimized LC-MS methods featuring run times of less than 1 minute, achieving the throughput required to process ≥10,000 samples per day presents significant challenges. This fundamental limitation creates a critical bottleneck that makes conventional chromatography incompatible with modern high-throughput screening (HTS) workflows. The innovative Acoustic Ejection Mass Spectrometry (AEMS) technology addresses these barriers by directly coupling acoustic droplet ejection with mass spectrometric detection, eliminating chromatographic separation and requiring only nanoliter-scale sample volumes.^11, 12^ This approach enables the precise measurement of targeted MRM transitions or acquisition of full-scan MS data from each well of 384-well or 1536-well microplates, supporting hundreds to thousands of measurements per hour. AEMS technology is a powerful tool for applications such as drug discovery, quality control, clinical diagnostics, and biomarker identification, helping to uncover disease mechanisms, drug mechanisms of action, and metabolic pathways.^13-20^

The advancement of high-throughput mass spectrometry has revealed significant computational limitations in data processing pipelines. AEMS instrumentation generates increasingly complex and voluminous datasets, yet the vendor-specific software solutions remain inadequate for large-scale studies due to four fundamental constraints: (1) lack of automated systems for tasks such as shot-to-peak alignment and multi-mode data integration (precursor ion (PI), and neutral loss (NL) scanning), (2) insufficient algorithms for heterogeneous data integration, (3) suboptimal processing throughput for high-density datasets, and (4) non-intuitive workflow configuration interfaces. While several open-source platforms (ASTS,^21^ MAVEN,^22^ MaxQuant,^23^ MRMAnalyzer,^24^ MRMPROBS,^25^ Skyline,^26^ SmartPeak,^27^ XCMS-MRM and METLIN-MRM,^28^ etc) provide essential functionality for data processing, sample segmentation, spectral feature extraction, and dataset aggregation, none currently address the specialized requirements of AEMS platforms, particularly their unique data architectures and ultra-high-throughput processing demands.

To bridge this gap, we developed rtmsEcho, an open-source R package extending our previously published rtms framework^29^. The rtmsEcho analytical pipeline comprises three core components (Figure 1): raw data file upload with associated plate/compound metadata; automated processing through optimized algorithms; and generation of standardized reports with interactive visualization tools for quality assessment. This streamlined workflow significantly improves both experimental reproducibility and analytical throughput, particularly for large-scale screening applications ranging from early-stage drug discovery to clinical translation research. By integrating robust automation with rigorous analytical standards, the system effectively addresses the growing demand for high-performance mass spectrometry data processing solutions capable of supporting modern high throughput.

**Figure 1.**
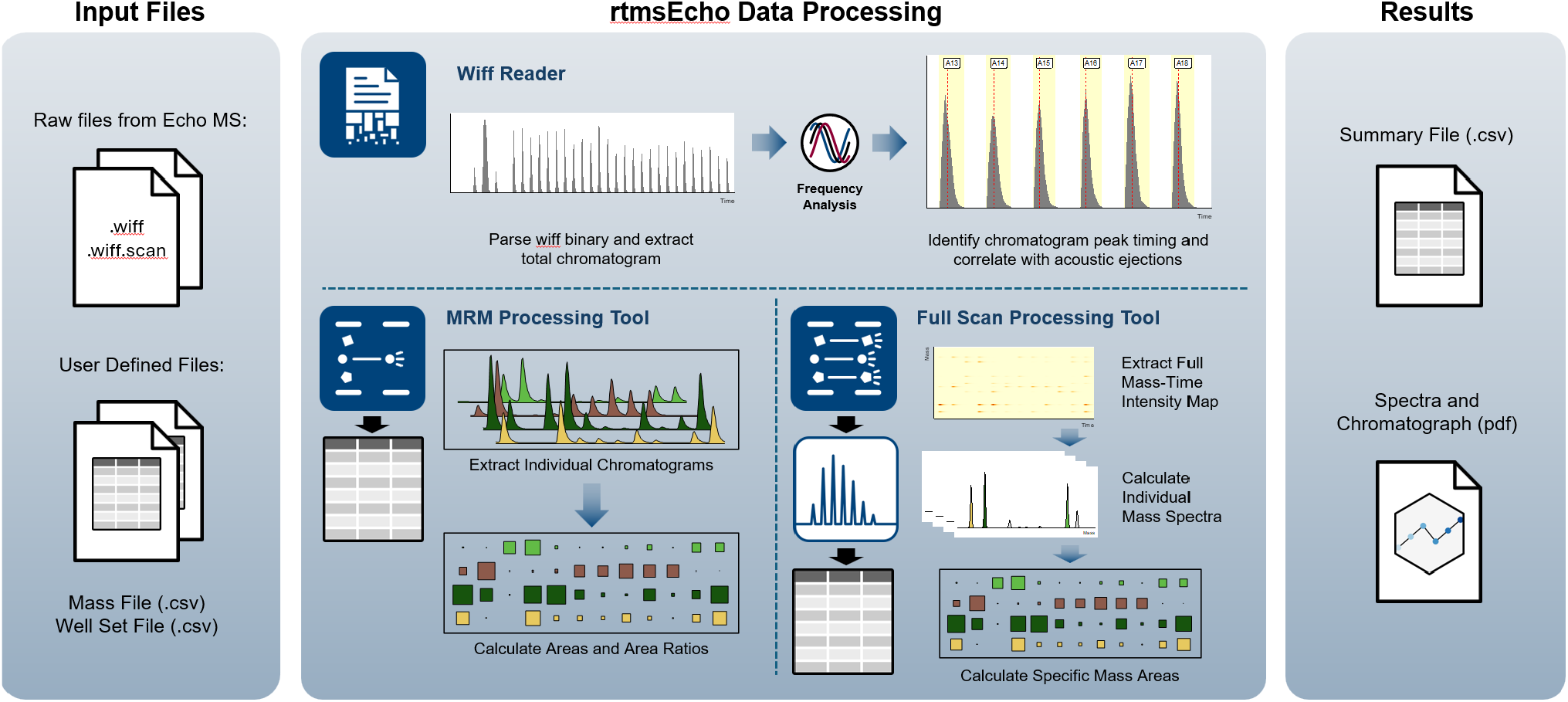
rtmsEcho Workflow. The rtmsEcho software enables fast, accurate, and automated processing of mass spectrometry data across targeted, semi-targeted, and untargeted mass spectrometry runs. a) rtmsEcho supports instrument data files in .wiff and .wiff.scan formats, along with user-defined fields containing sample and analyte information. b) rtmsEcho integrates underlying algorithms by parsing the .wiff binary data and extracting the total chromatogram from Microsoft’s Compound File Binary Format. Basic Fourier analysis is used to identify the maximum frequency and phase where the total ion chromatogram signal best aligns with a continuously varying sinusoid across the entire run. Peaks from this sine wave are then assigned, and the measured ejections of the run are marked. For MRM, individual transitions are extracted from the TIC. Furthermore, the peak area of each transition and the area ratio of the analyte to the assigned internal standard were calculated. In the full scan processing section for untargeted runs (precursor ion and neutral loss), mass-time intensity maps are extracted from the .wiff file, individual mass spectra are calculated, and mass areas are determined for each mass ion. c) A reporting configuration, chromatograph review, and summary files are provided after executing the data processing workflow, allowing results to be reviewed and validated before final reporting.

### EXPERIMENTAL SECTION

#### MRM Data Acquisition Mode

MRM was used for the quantification of target analytes via the two-stage mass filtering process: precursor ion selection and fragmentation followed by specific product ion monitoring (Figure 2a). Using a commercial screening library (APEXBIO, 10 mM in DMSO), we conducted three complementary experiments: (1) Preparation of analytical working solutions, where eight compounds (Cytosine-1-β-D-arabinofuranoside hydrochloride, Gemcitabine HCl, Lamivudine, Zalcitabine, Flubendazole, Mebendazole, Albendazole, and Fenbendazole) were diluted with DMSO to generate concentration series (up to 1 μM) in 384-well plates; (2) Quantitative analysis of carbamazepine and warfarin in quality control plates using serial dilutions according to an established protocol;^16^ and (3) Metabolic profiling of library compounds, as described previously ^30^. These comprehensive datasets were used to optimize the software algorithms and validate the reliability and robustness of the data processing pipeline.

**Figure 2.**
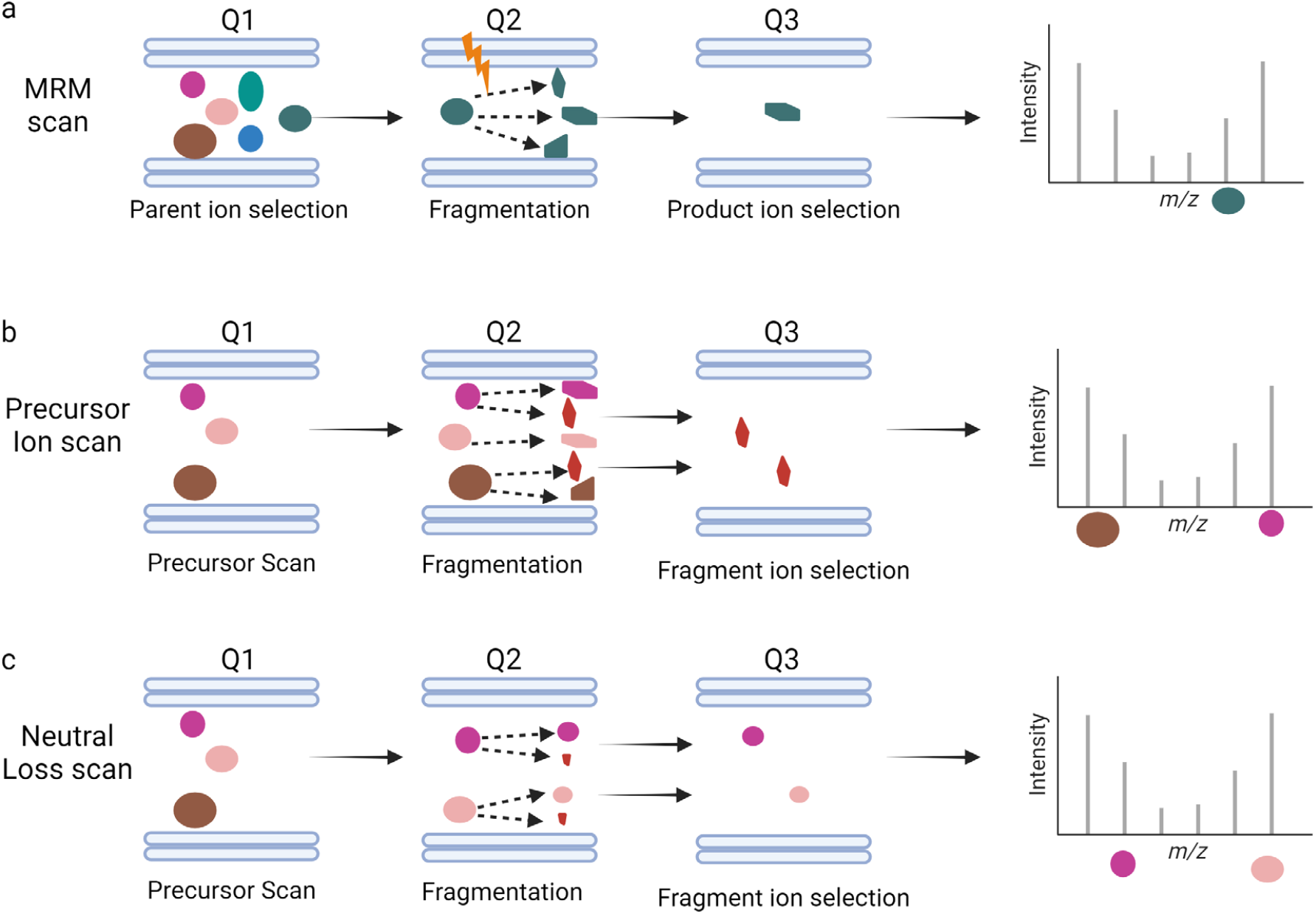
Quantitative scan modes. a) MRM Scan Mode: In this mode, one or more specific ion transitions are selected, with the parent ion being detected in Q1 and the product ion in Q3. Only ions that match the exact parent-product ion pair are detected, ensuring high sensitivity and accurate quantification of the compound. b) Precursor Ion Scan Mode: This mode scans all ions within a set mass range in Q1 and filters them for a specific fragmental ion in Q3, allowing for the detection of precursor ions. c) Neutral Loss Scan Mode: In Q1, all ions within a specified mass range are scanned, and in Q3, only the fragment ions that lost a specific neutral fragment are selected for detection.

#### Full Scan Data Acquisition Mode

The complementary NL and PI scanning modes provide analytical specificity for complex molecules by targeting either neutral fragments (NL mode) or diagnostic precursor ions (PI mode) (Figures 2b, 2c). These techniques deliver the sensitivity and selectivity required for advanced applications in quantitative analyses^17^. For PI mode analysis, we selected four nucleoside analogs (Cytosine-1-β-D-arabinofuranoside hydrochloride, Gemcitabine HCl, Lamivudine, and Zalcitabine; Figure 3a) based on their identical product ion (m/z 112). Similarly, four benzimidazole compounds (Flubendazole, Mebendazole, Albendazole, and Fenbendazole; Figure 3b) were chosen for NL analysis due to their common loss of a methanol group (CH3OH, 32 Da). All samples were prepared following the protocol described above.

**Figure 3.**
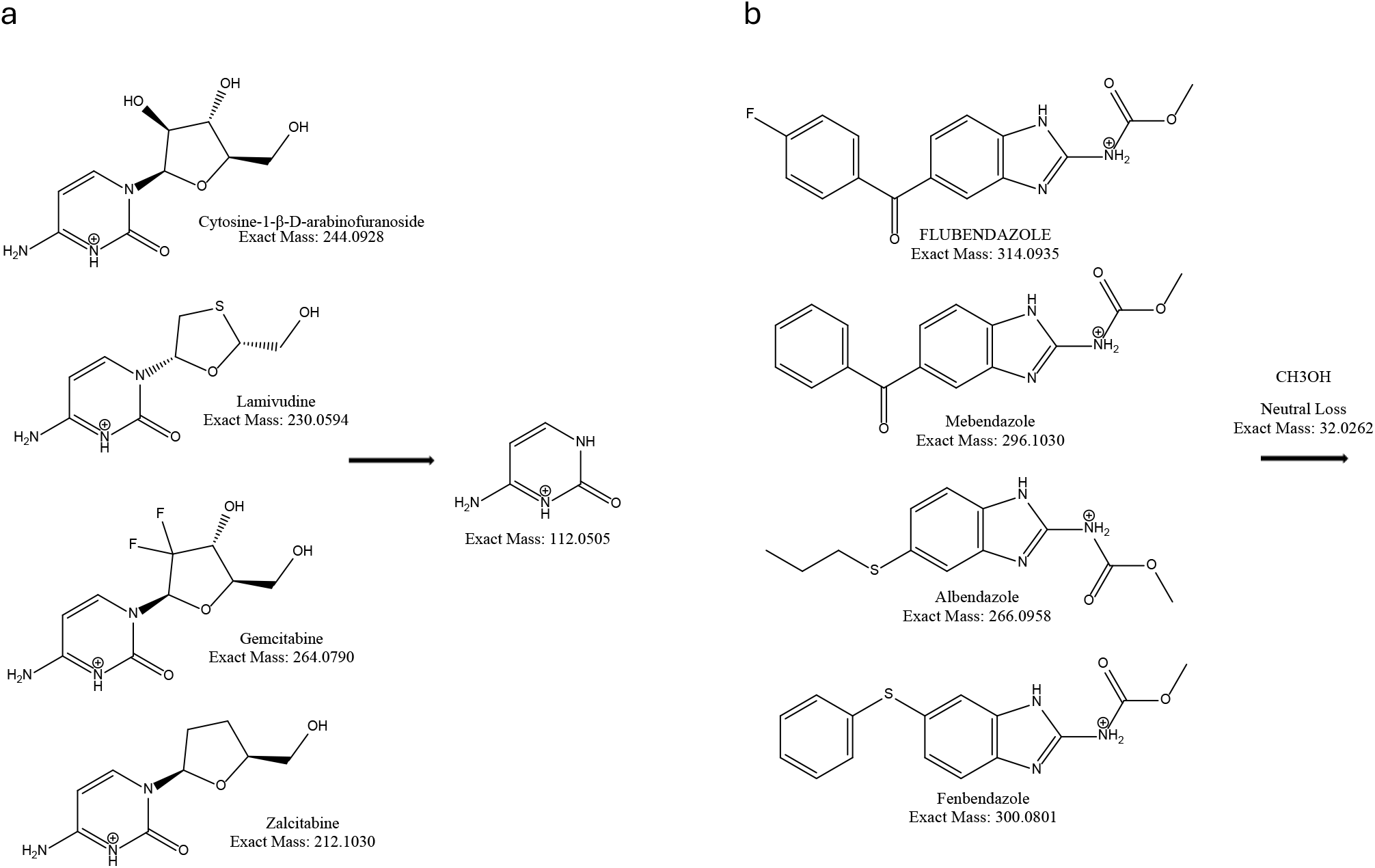
Molecule structures selected for testing rtmsEcho. a) Four molecules were chosen for both MRM and precursor ion scans, all having the same product ion of 112. b) Four molecules were selected for both MRM and neutral loss scans, all exhibiting the same ion loss of 32.

#### AEMS system

All mass spectrometric analyses were performed using an AB Sciex Triple Quad 6500+ system (Sciex, Concord, ON, Canada) equipped with an AEMS acoustic autosampler and controlled by Sciex OS Analytics Software (v3.1.0). The system was operated in both multiple reaction monitoring (MRM) or full-scan acquisition modes with positive ion electrospray ionization (ESI). A methanol-based mobile phase containing 1 mM ammonium fluoride was delivered at 400 µL/min as the carrier solvent. Samples (up to 20 nL) were acoustically ejected from 384-well polypropylene plates into the open port interface (OPI) and subsequently introduced into the OptiFlow Turbo V ion source through a transfer capillary. The acoustic ejection was performed in normal mode with a 1000 ms interval and 2000 ms method delay. The ESI source parameters were optimized as follows: nebulizer gas (GS1) 90 psi, heater gas (GS2) 70 psi, curtain gas 20 psi, collision gas (CAD) 9 units, source temperature 500°C, and ion spray voltage 5500V. Data acquisition employed a 15 ms dwell time with 5 ms pause between transitions. The .wiff files from runs in MRM mode were processed with Sciex OS, peaks were integrated using the following criteria: minimum signal-to-noise ratio (S/N) of 2, minimum peak height of 100 counts, and expected retention time window of ≥0.02 minutes. However, as Sciex OS could not process the PI and NL mode data, the raw time, area dataset was exported from the Sciex OS software, then manually aligned with the correct ejection time and assigned the area of that ejection.

### rtmsEcho Workflow

#### Data Representation

Every AEMS .wiff file contains a data structure that we refer to as the total ion chromatogram (TIC). This data structure contains a single intensity for every detection time point, representing the total intensity measured by the detector at that moment, whether it aggregates data from either multiple explicit mass transitions in MRM mode or across all values in an m/z range in NL or PI mode. The TIC in the .wiff file also contains binary data necessary to locate the particular measurements for that detection time point in the .wiff.scan file, as well as a flag distinguishing between a discrete set of several intensities (MRM mode) or a compressed full spectra (PI or NL mode).

In the .wiff.scan file, at the specified file offset, the more detailed intensities collected at a particular detection time point are stored. For MRM files, this is simply a set of intensity values, one for each mass transition specified (Figure 4A, 4C, and 4E); in the case of PI or NL runs, it is a compressed binary data structure encoding a full array of intensity values, and their exact *m/z* values (Figure 4D, and 4F).

**Figure 4.**
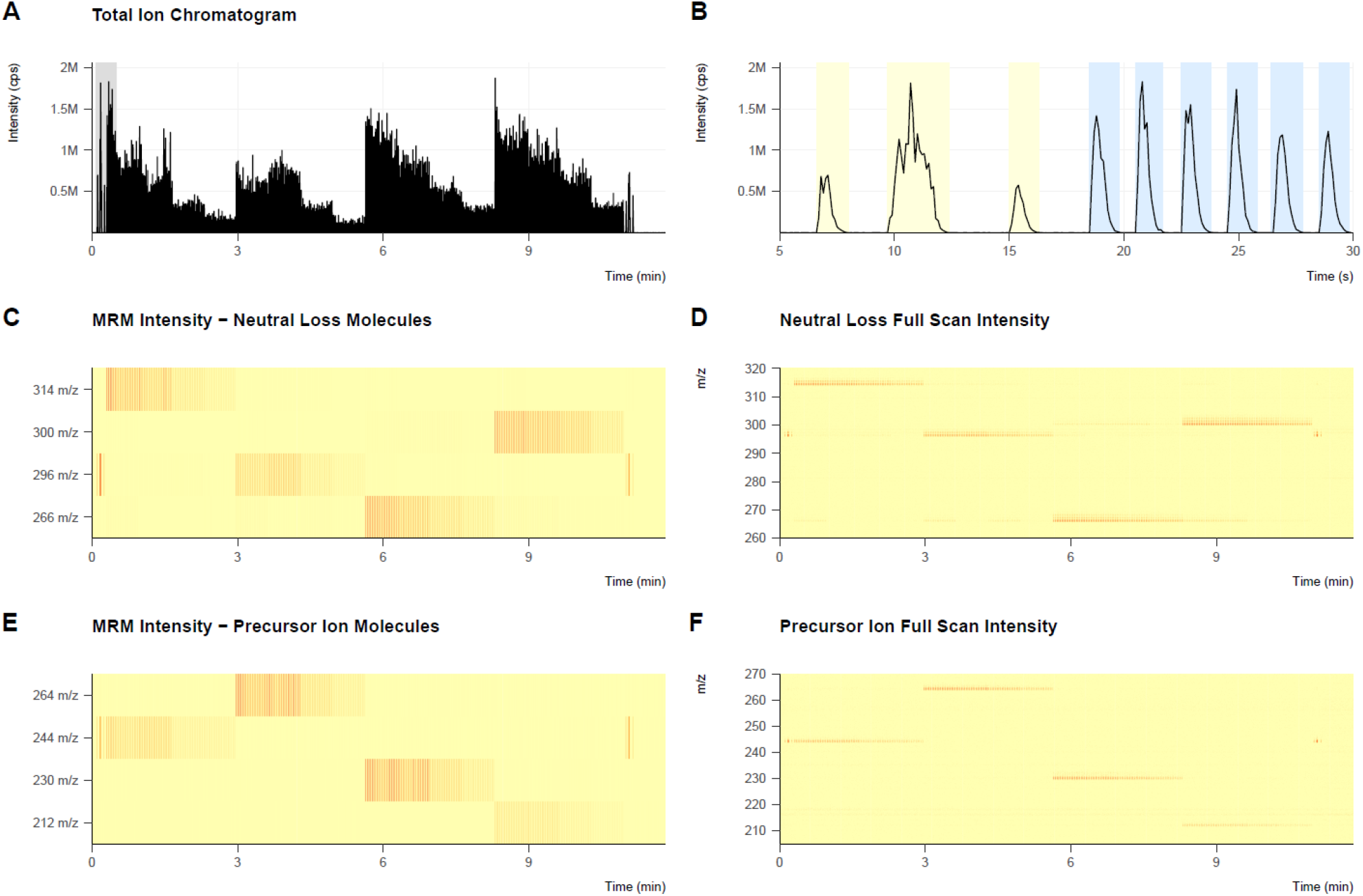
Representative Chromatogram profiles. (A) Plot of the total ion chromatogram for a single Echo MS run. The plot shows the total ion intensity across all masses measured at each detection time point across the entire run. The area highlighted in gray is depicted in figure B. (B) Zoomed in slice of the total ion chromatogram from figure A, ranging from 5 seconds into the run to 30 seconds into the run. The short-long-short barcode used by Sciex OS to analyze the signal can be seen highlighted in yellow, whereas the first six measurement ejections of the run can be seen, bounded in blue. These latter six times and bounds are automatically extracted by rtmsEcho. (C) The intensity of four explicit mass transitions measured at each detection time point during an MRM run. This run was collected using mass transitions to identify the same masses and ions as those detected in the neutral loss run depicted in figure D. (D) Intensities of all measured m/z values at all detection time points across an entire neutral loss run. This run was performed using the same compounds, plate, and timing as the MRM data depicted in figure C. (E) The intensity of four explicit mass transitions measured at each detection time point during an MRM run. This run was collected using mass transitions to identify the same masses and ions as those detected in the precursor ion run depicted in figure F. (F) Intensities of all measured m/z values at all detection time points across an entire precursor ion run. This run was performed using the same compounds, plate, and timing as the MRM data depicted in figure E.

#### Data quantification

The precise time in a run at which each ejection occurs and is detected is not explicitly stored in the .wiff file or the .wiff.scan file. Instead, the timing of the ejections must be inferred from the structure of the data. rtmsEcho infers the timing of ejections as a complete set based on the assumption that acoustic ejections – and therefore peaks in intensity resulting from those ejections – will occur at evenly spaced times across the run. With a standard estimate of the start time of the run and a fairly precise initial estimate of the number of milliseconds between successive peaks, optimizing the agreement between the total intensity signal over time and a simple sinusoidal signal can identify the true timing of peaks to within 0.01 milliseconds. Optimizing this sine wave sets the precise time of the expected max intensity for all ejections in a run.

To determine the range of times during which an ejection is being detected, rtmsEcho performs a slight smoothing of the TIC intensity signal and locates the local minima between adjacent peak times. The beginning and end of a window of time during which an ejection is determined to be detected is then either the range of time during which the total intensity is above some threshold of background noise, or, if the signal never drops below the background threshold between adjacent ejections, the local minimum in smoothed total intensity. At the end of these calculations, rtmsEcho has estimated a peak time, minimum time, and maximum time for the detection of each ejection in the run.

##### Quantification Details – MRM

As the data in the .wiff.scan file for MRM runs is already explicitly broken into several discrete mass transitions, it can be directly extracted as a discrete set of XICs (Figure 4). Therefore, once the timing of ejections is estimated, the quantification of a particular mass for each ejection is simply a matter of taking the total intensity of the XIC for that mass transition in the window of time associated with that ejection. To account for the presence of baseline background intensity for a given mass transition, rtmsEcho uses a consistent baseline correction for a given mass transition across an entire run. The baseline intensity for a given mass transition is estimated by taking the average intensity for that transition in two one-second windows, before the first ejection and after the last ejection. To account for the fact this baseline will overlap with measured intensities rather than simply be added to them, it is scaled by a factor of 2/3, and then all subtracted from all intensities in the run. Those that lie below zero are set to zero, and the quantified peak area for a given ejection is the sum of these lowered intensities.

##### Quantification Details – Full Scan

In PI or NL modes, the data stored in the .wiff.scan file for a given detection time point is a full mass spectrum representing all intensities measured across all m/z values. The total resulting data structure is therefore a massive array of intensities, one for each detection time point *and* m/z value. To quantify the presence of a particular mass in a particular ejection, this array must be narrowed both to the correct range of masses and the correct range of times. In rtmsEcho, given a range of detection time points estimated from the TIC in the .wiff file, all intensities for a given m/z value are integrated in that range, producing a total intensity measured for each m/z value in that ejection. This spectrum is a data structure compatible with the previously published rtms R package ^29, 31^ and can be analyzed, quantified, and plotted using the methods of that package. This method leverages the more stable ejection timing determined by the TIC and takes full advantage of the full scan approach, allowing the user to quantify unanticipated mass peaks that arise from a run. The rtmsEcho full-scan methods therefore support both targeted and untargeted mass quantification.

##### R package

Development of the package was performed in R, an open-source programming and data science environment ^32, 33^, and the RStudio IDE ^34^. The package was developed using the devtools package suite ^35^, and for comparative analyses between Sciex OS and rtmsEcho processed data, all data were analyzed in R and RStudio and visualized using the ggplot2 package ^36^.

### RESULTS AND DISCUSSION

#### Comparison of Sciex OS and rtmsEcho

##### MRM Quantification

In MRM-based AEMS analysis, the TIC represents the composite signal of all monitored mass transitions. As illustrated in Figure 4A, distinct compounds (four analytes injected at 80-ejection intervals each) generate characteristic intensity profiles within the TIC (Figure 4C). Traditional processing via Sciex OS software quantifies these transitions by aligning ejection events using a critical but fragile “barcode” signal, a short-long-short ejection pattern from a designated well. Failure of any barcode component renders the dataset unusable, requiring reruns. In contrast, our open-source rtmsEcho package circumvents this limitation by employing TIC-derived peak detection to define ejection windows, eliminating dependency on barcode signals. Validation against Sciex OS (Figure 5A) demonstrated exceptional concordance (R^2^ = 100%) for 640 samples across two 384-well plates, with intensities spanning 0–800,000 counts. Notably, rtmsEcho achieves this while automating transition assignments—a manual step in vendor software—and accelerating processing by >10-fold through optimized algorithms.

**Figure 5.**
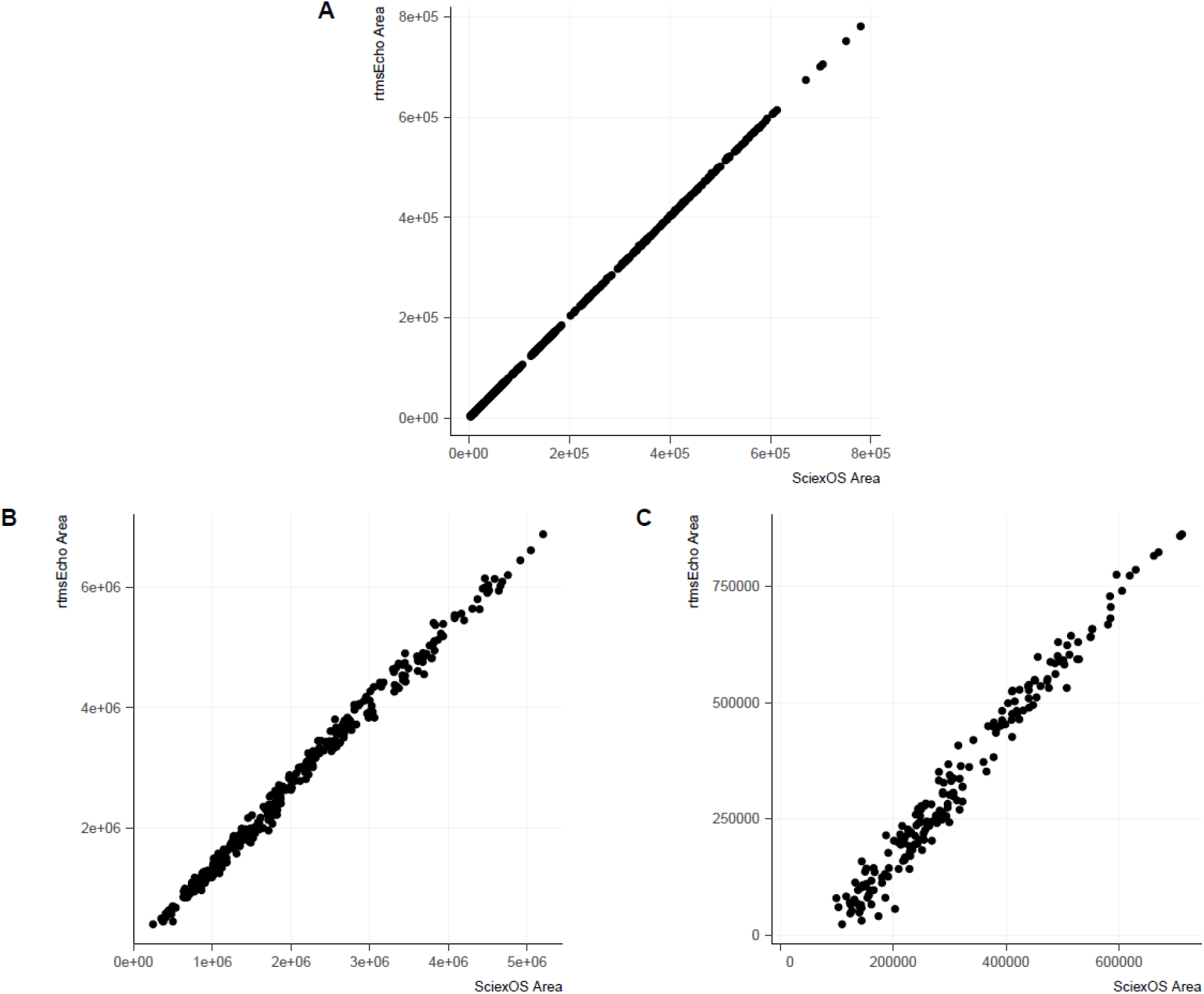
Comparison of peak area quantification between SCIEX OS and rtmsEcho: a) Data acquired using MRM scan mode demonstrate a strong alignment between area quantification by SCIEX OS and rtmsEcho (r^2^ = 1, n = 640). b) Data acquired using precursor ion mode show good agreement in area quantification between manual alignment from SCIEX OS and rtmsEcho (r^2^ = 0.9744, n = 320). c) Data acquired using neutral loss mode reveal excellent alignment in area quantification by manual alignment from SCIEX OS and rtmsEcho (r^2^ = 0.9927, n = 320).

The asynchrony between acoustic ejections and mass spectral acquisition in AEMS makes it difficult to properly align ejections to the chromatographic peak, particularly in noisy or full-scan modes. While Sciex OS is capable of processing MRM data (targeting predefined transitions), it is incapable of analyzing untargeted scan modes. rtmsEcho addresses these gaps by providing direct access to raw instrument data, enabling experiment-specific workflows. For instance, it supports dynamic transition mapping without manual intervention and accommodates exploratory analyses beyond the targeted scope of MRM. This flexibility, combined with robust peak integration and baseline correction, positions rtmsEcho as a versatile solution for high-throughput AEMS datasets.

##### Full-scan Data

Full-scan acquisition modes (PI and NL scans) in AEMS present both unique opportunities and significant analytical challenges compared to traditional MRM approaches. Unlike targeted MRM runs that measure predefined mass transitions, full-scan modes capture complete mass spectra at each time point across a specified m/z range (Figure 4D, 4F). This comprehensive detection capability enables retrospective analysis of unexpected compounds and metabolites, making it particularly valuable for untargeted screening applications. However, the resulting datasets are substantially more complex, often containing noisy baseline signals that can obscure analyte peaks (Figure 4D, 4F). These challenges are compounded by the lack of software available to process full-scan data from AEMS instruments.

The rtmsEcho package addresses these limitations through innovative data processing strategies. When analyzing full-scan data with persistent high-intensity baseline signals, rtmsEcho can selectively use well-defined high-mass peaks (Figure 4D) as reference points for peak alignment, subsequently applying these time frames to quantify less well-defined, lower signals at other m/z values. This approach provides efficient processing, reducing analysis times of a 384-well plate from 3 hours (with manual alignment from the data profile) to just 30 seconds with rtmsEcho -a 360-fold improvement while maintaining excellent quantification agreement (R^2^ > 0.97) between methods. Systematic differences in absolute area measurements were observed between the platforms, with Sciex OS typically reporting values 15-20% lower than rtmsEcho (Figure 5B). This observed discrepancy originates from distinct baseline correction methodologies: Sciex OS utilizes a local minima subtraction algorithm, whereas rtmsEcho employs a global noise floor estimation technique. The local minima approach tends to overestimate baseline levels in high-intensity regions due to its restricted detection window, while the global method establishes a more consistent baseline reference across the entire chromatogram. Consequently, when analyzing compounds with similar signal magnitudes (whether targeted or untargeted), rtmsEcho’s approach yields higher net peak intensities as it accounts for instrumental background noise.

Similar performance gains were observed for NL scans, where processing time decreased from 1 hour to 30 seconds while maintaining excellent quantification agreement (R^2^ > 0.99) between methods (Figure 5C). The quick processing speed of rtmsEcho allows for large-scale validation studies that would be impractical with conventional software, facilitating robust method development and quality control for full-scan AEMS applications.

These advancements position rtmsEcho as an essential tool for unlocking the full potential of AEMS instruments in full-scan modes. By overcoming the processing limitations of vendor software while delivering superior speed and flexibility, rtmsEcho enables researchers to harness the rich information content of full-scan AEMS data for both targeted and untargeted analyses. Future developments will focus on expanding these capabilities to additional scan modes and implementing advanced machine learning algorithms for automated peak detection and quantification.

#### Validation of rtmsEcho for Large-Scale Compound Screening

##### QC plates Validation

To rigorously evaluate the quantification accuracy or rtmsEcho, we analyzed data from 20 quality control plates acquired in MRM mode. This validation focused on two key aspects: (1) raw quantification agreement between rtmsEcho and Sciex OS, and (2) precision in concentration-dependent responses.

For the first test, we selected two representative compounds—warfarin and carbamazepine—and compared their estimated peak areas between methods. The results demonstrated near-perfect alignment, with R^2^ values virtually indistinguishable from 1.0 (0.9998 for warfarin, which served as the internal standard, and 0.9975 for the area ratio of carbamazepine to warfarin; see Table 1, Supplemental Figures 1a and 1b), confirming that rtmsEcho replicates the data processing and quantification of Sciex OS.

**Table 1.** The agreement between SciexOS outputs and rtmsEcho outputs for MRM datasets.

|  | # of MRM transitions | # of data points | R<sup>2</sup> |
| --- | --- | --- | --- |
| *QC Plate* |  |  |  |
| Internal Standard (Warfarin) | 1 | 7680 | 0.9998 |
| Area Ratio (Carbamazepine) | 2 | 7680 | 0.9975 |
| *Biochemical Assay* |  |  |  |
| Internal Standard (Warfarin) | 1 | 17856 | 0.9974 |
| Area Ratio (All Targets) | 1489 | 17856 | 0.9895 |

To assess concentration-dependent accuracy, we examined carbamazepine area ratios (analyte/internal standard) plotted against known concentrations on a logarithmic scale. Both rtmsEcho and Sciex OS produced superimposable correlation trends (R^2^ = 0.995; see Supplemental Figures 1c and 1d), with comparable variability across three orders of magnitude. This demonstrates that rtmsEcho preserves the precision of the vendor software, Sciex OS, in quantifying relative abundances—a critical requirement for assays relying on area ratios, such as pharmacokinetic studies or enzyme activity screens.

#### Large-Scale CYP450 Assay Validation

We conducted an extensive evaluation of the performance of rtmsEcho in analyzing a high-throughput CYP3A4/5 substrate assay comprising 1,489 validated compounds^30^. This dataset included measurements of the internal standard, warfarin, and area ratios of target compounds to warfarin. Despite the complexity of the dataset, both rtmsEcho and Sciex OS demonstrated strong linearity in their area ratio calculations, with R^2^ values of 0.9974 for warfarin peak area alone and 0.9895 for the area ratios across 17,856 data points.

Internal standard correlations reflect the squared correlation between the peak area reported for warfarin by Sciex and the corresponding area reported by rtmsEcho across all points in a dataset, calculated on a linear scale. Area ratio correlations reflect the squared correlations between area ratios calculated for the MRM mass present in each well, as reported by SciexOS and rtmsEcho, calculated on a logarithmic scale.

These results underscore the robustness and reliability of rtmsEcho in high-throughput biochemical screening and drug interaction studies. The high-throughput capabilities of rtmsEcho, coupled with its accuracy and precision, make it a powerful tool for large-scale drug discovery studies. The ability to process thousands of samples rapidly without compromising data quality is crucial for the timely assessment of high throughput screening and quantitative analysis. Moreover, the strong linearity observed across a broad range of compounds highlights the versatility of rtmsEcho and the potential for integration with diverse analytical workflows.

#### Throughput and Usability Advantages

Beyond accuracy, rtmsEcho offers practical benefits for large datasets: 1) The elimination of manual transition mapping: Unlike Sciex OS, which requires predefined transition lists, rtmsEcho auto-assigns transitions using metadata, reducing errors; 2) Faster processing: rtmsEcho R code cuts analysis time by more than 90% for plates with 384 or 1536 wells (benchmark data not shown); and 3) Native integration with R ecosystems: Enables direct piping of results into statistical models (e.g., dose-response fitting with dose response) or visualization tools (e.g., ggplot2)^36^. These features make rtmsEcho particularly suited for large-scale screens, where reproducibility and pipeline automation are paramount.

## CONCLUSIONS

In summary, rtmsEcho represents a significant advancement in the processing and analysis of Acoustic Ejection Mass Spectrometry data, providing a robust, open-source solution that bridges the gap between high-throughput experimentation and efficient data interpretation. Developed as an R package, rtmsEcho supports multiple scan modes—including MRM, PI, and NL modes—delivering unparalleled speed and accuracy in data processing. By automating critical steps such as peak alignment, baseline correction, and area quantification, the package reduces processing times from days to minutes, achieving efficiency gains of 10-to 100-fold compared to conventional methods. This dramatic acceleration does not come at the expense of data quality; instead, rtmsEcho enhances reproducibility and precision, ensuring reliable results for both targeted quantitation and untargeted screening applications.

Beyond its computational efficiency, rtmsEcho offers great flexibility, allowing researchers to customize analyses to meet the specific demands of their experiments. Its integration with the R ecosystem enables advanced downstream statistical and graphical analyses, further extending its use in high-throughput environments. By eliminating the need for manual transition mapping and proprietary software dependencies, rtmsEcho increases access to high-performance AEMS data processing, making it a valuable tool for academic, pharmaceutical, and industrial laboratories. Future developments will focus on expanding its functionality to broader compatibility with additional mass spectrometry platforms. Ultimately, rtmsEcho sets a new benchmark for rapid, automated, and reproducible mass spectrometry data analysis.

## AUTHOR INFORMATION

### Corresponding Author

**Lei Yang-** Analytical Technologies Center, Department of Chemical Biology and Therapeutics, St. Jude Children’s Research Hospital, Memphis, Tennessee;

### Authors

**Mary Ashley Rimmer**-Analytical Technologies Center, Department of Chemical Biology and Therapeutics, St. Jude Children’s Research Hospital, Memphis, Tennessee;

**Nathaniel Twarog**-Lead Discovery Informatics, Department of Chemical Biology and Therapeutics, St. Jude Children’s Research Hospital, Memphis, Tennessee;

**Tharindu Ranathunge**-Analytical Technologies Center, Department of Chemical Biology and Therapeutics, St. Jude Children’s Research Hospital, Memphis, Tennessee;

**Jingheng Wang**-Department of Chemical Biology and Therapeutics, St. Jude Children’s Research Hospital, Memphis, Tennessee;

**Yong Li**-Analytical Technologies Center, Department of Chemical Biology and Therapeutics, St. Jude Children’s Research Hospital, Memphis, Tennessee;

**Taosheng Chen**-Department of Chemical Biology and Therapeutics, St. Jude Children’s Research Hospital, Memphis, Tennessee;

**Anang Shelt**-Lead Discovery Informatics, Department of Chemical Biology and Therapeutics, St. Jude Children’s Research Hospital, Memphis, Tennessee;

### Author Contributions

M.R., N.T., wrote the R algorithms, wrote the rtmsEcho application, and wrote the manuscript. L.Y. designed and performed the experiments and wrote the manuscript. Y.L., T.R., and J.W. conducted AEMS experiments, processed the data, and reviewed the manuscript. T.C. and A.S. reviewed the manuscript.

### Notes

The authors state that they have no competing financial interests or personal relationships that may have influenced the work presented in this manuscript.

## ACKNOWLEDGMENTS

We are grateful for the support of the American Lebanese Syrian Associated Charities (ALSAC) and St. Jude Children’s Research Hospital, and we would like to thank the patients, their families, and the staff at our institution. We acknowledge the support and contribution of all the personnel from the Department of Chemical Biology and Therapeutics at St. Jude who were involved in the earlier R&D stages of this work and in supplying reagents.

## Supplemental materials

**Supplemental Figure 1.**
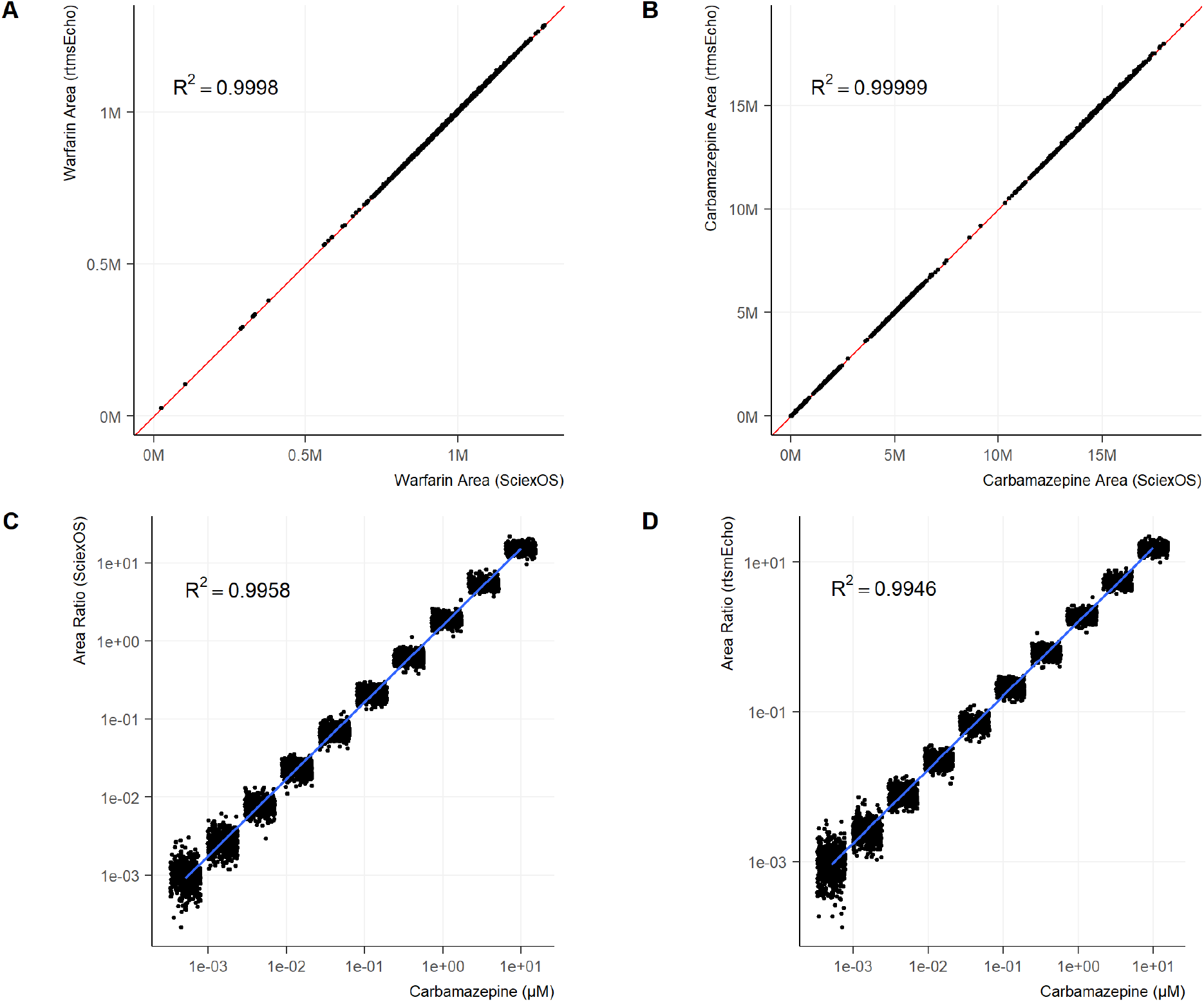
20 Compound Management QC plates were analyzed using MRM scanning mode. (A) For raw quantification, Sciex OS and rtmsEcho produce nearly identical results for the compounds warfarin and carbamazepine. The estimated areas for both compounds show a correlation between methods with an R-squared value virtually indistinguishable from 1. (B, C) When comparing the area ratios for carbamazepine estimated by Sciex OS and rtmsEcho to the actual carbamazepine concentrations on a logarithmic scale, both methods demonstrate nearly identical precision in their correlation.

**Supplemental Figure 2.**
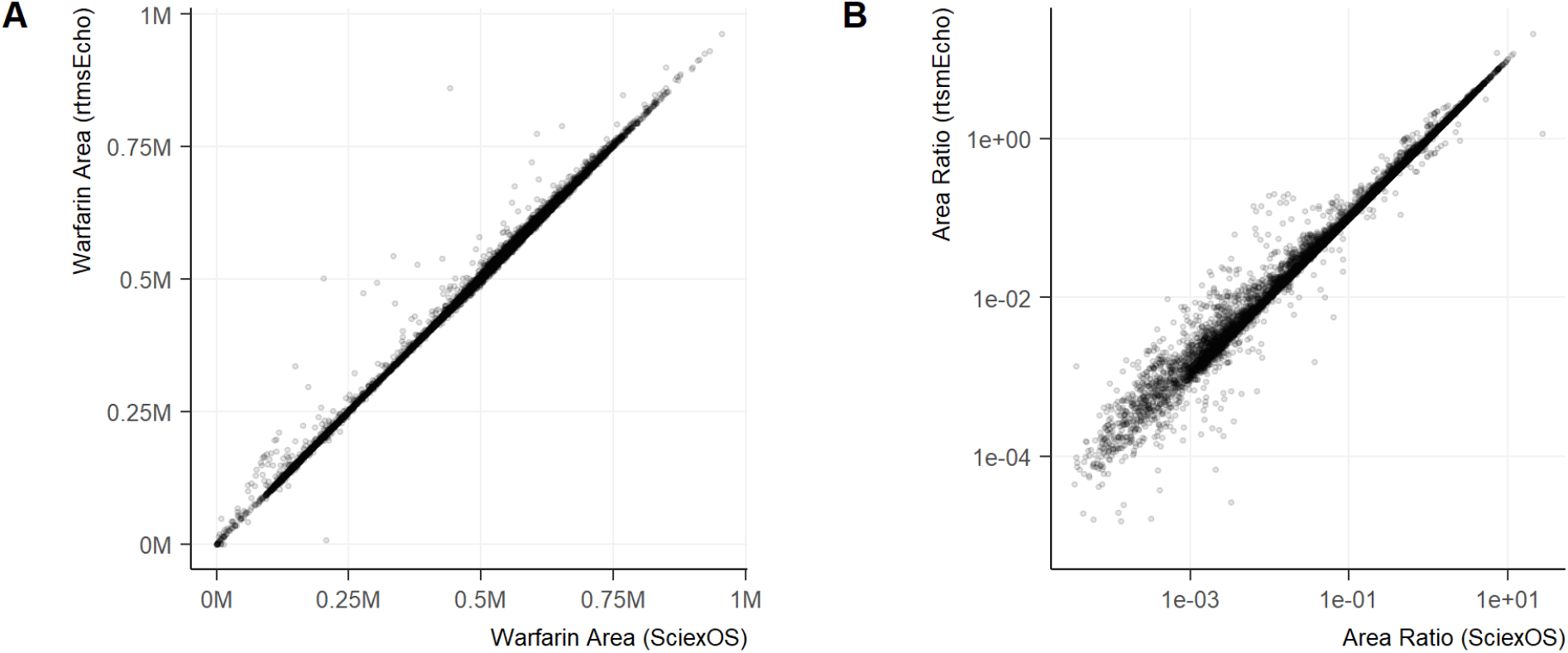
A CYP 3A4/5 selective assay using MRM scan for 1500 compounds shows that the quantification of area ratios by Sciex OS and rtmsEcho remain closely aligned, with R-squared values of 0.998 and 0.977 on linear and logarithmic scales, respectively.

